# Magnesium ions mediate ligand binding and conformational transition of the SAM/SAH riboswitch

**DOI:** 10.1101/2023.03.12.532287

**Authors:** Guodong Hu, Huan-Xiang Zhou

## Abstract

The SAM/SAH riboswitch binds S-adenosylmethionine (SAM) and S-adenosylhomocysteine (SAH) with similar affinities. Mg^2+^ is generally known to stabilize RNA structures by neutralizing phosphates, but how it contributes to ligand binding and conformational transition is understudied. Here, extensive molecular dynamics simulations (totaling 120 μs) identified over 10 inner-shell Mg^2+^ ions in the SAM/SAH riboswitch. Six of them line the two sides of a groove to widen it and thereby pre-organize the riboswitch for ligand entry. They also form outer-shell coordination with the ligands and stabilize an RNA-ligand hydrogen bond, which effectively diminish the selectivity between SAM and SAH. One Mg^2+^ ion unique to the apo form maintains the Shine-Dalgarno sequence in an autonomous mode and thereby facilitates its release for ribosome binding. Mg^2+^ thus plays vital roles in SAM/SAH riboswitch function.

## INTRODUCTION

Although RNAs are best known for transferring genetic information from DNA to proteins, some RNAs such as riboswitches perform signaling and catalytic functions much like proteins ^1^. Riboswitches, mostly found in bacteria, are located in the 5’-untranslated regions of mRNAs and typically consist of two domains: aptamer and expression platform ^2, 3^. The aptamer domain binds ligands such as metabolites and triggers the expression platform to turn on or off gene expression ^4, 5^. S-adenosylmethionine (SAM; 0A) is the major methyl donor for the methylation of nucleic acids and proteins and is thus an essential metabolite ^6–9^. It consists of an aminocarboxypropyl, a positively charged sulfonium center substituted by methyl, and a 5’-deoxyadenosyl. After donating its methyl, SAM is converted to the neutral compound S-adenosylhomocysteine (SAH; 0A). Based on structure, sequence, and evolutionary relatedness, SAM-sensing riboswitches have been divided into several families ^10, 11^, including SAM-I ^12, 13^, SAM-II ^14^, SAM-III ^15^, SAM-IV ^16^, and SAM-I/IV ^11^, SAM-V ^17^, SAM-VI ^18, 19^, and SAM/SAH ^11^, ^20^. SAM/SAH riboswitches are distinct by their similar affinities for binding SAM and SAH, and by their smaller size and lower complexity of the ligand-binding pocket ^11, 20^.

A high-resolution NMR structure of the env9b SAM/SAH riboswitch bound with SAH was recently determined by Weickhmann *et al*. (Protein Data Bank entry 6HAG) ^20^. The structure features an H-type pseudoknot (0B, C), which consists of two stems: A-form stem S1 formed by pairing the G1 to C5 bases with the C24 to G20 bases, and pseudoknotted stem S2 with five canonical pairs between G10, C11, U12, C14, and C15 in the 5’ strand and C42, G41, A40, G39, and G38 in the 3’ strand. Two short loops, L1 (A6 to G9) and L2 (U16 to C19), connect the S1 and S2 stems; a third long, flexible loop, L3 (A25 to U37), connects the 3’ end of S1 to the 5’ end of S2. A sandwich-shaped ligand-binding pocket is formed between the C8:G17 base pair and the C15:G38 base pair (0C). The nucleobases lining the ligand-binding pocket mostly interact with the 5’-deoxyadenosyl group of the ligands. This group is stacked between G38 on the ceiling and C8:G17 on the floor. The sugar portion of the 5’-deoxyadenosyl group forms both CH/π interactions with the base of G9 on the ceiling and hydrogen bonds with the phosphate-sugar backbone of G9. The base portion of the 5’-deoxyadenosyl group forms a reversed Hoogsteen base pair with the U16 base. Recent crystal structures of the SK209-2-6 SAM/SAH riboswitch (lacking the highly flexible nucleotides 26-34 of L3) bound with SAM or SAH are very similar to the NMR structure of the SAH-bound env9b SAM/SAH riboswitch ^21^.

At the 3’ end of the riboswitch, the Shine-Dalgarno (SD) sequence (G38-G39-A40-G41) can be freed to bind with the ribosome and initiate translation when no ligand is present ^11, 20, 21^. As just noted, G38 base-stacks with the ligand. G39 forms a base triple with two other nucleobases, G9 and C14 (0C). As G9 interacts with the ligand, it serves as a bridge between the ligand and G39. Via these direct and indirect interactions, the ligand sequesters the SD sequence and maintains the riboswitch in the translational off state. In the apo form of the env9b riboswitch, broad peaks in the imino proton NMR spectrum suggested that the riboswitch is only partially structured and conformationally heterogeneous ^20^. Likewise, for the SK209-2-6 riboswitch, in-line probe indicated that nucleotides that form the pseudoknotted stem become less accessible upon SAM binding ^11^; single-molecule FRET revealed that the apo form is in dynamic exchange between partially and fully folded conformations ^21^.

Because electrostatic repulsion of the phosphate backbone of RNAs would lead to unstable tertiary structures, cations are essential to provide charge neutralization ^22^. Due to its small size and double charge, Mg^2+^ is special for RNA structural stability, e.g., by forming bidentate coordination with two adjacent phosphates ^23^. Moreover, Mg^2+^ has been implicated in many other roles, including promoting folding or conformational transition ^24–26^ or rescuing misfolding ^27^, mediating ligand binding ^20, 28–34^, and participating in catalysis ^35^. In particular, although imino proton NMR spectra demonstrated that the env9b SAM/SAH riboswitch is capable of binding SAH in the absence of Mg^2+^, isothermal titration calorimetry measurements showed that Mg^2+^ significantly increases the binding affinities of the riboswitch for both SAH and SAM ^20^. However, identifying Mg^2+^ ions in RNA structures by experimental approaches is very challenging. Although the diamagnetic Mg^2+^ in theory can lead to changes in NMR spectra of RNA, the effects may be small and challenging to resolve ^36^. For example, the NMR structure 6HAG, though determined on samples in 2 mM Mg(OAc)_2_, has no information on Mg^2+^. For X-ray diffraction, because Mg^2+^, Na^+^, and water molecule all have the same number of electrons, distinguishing them is difficult and requires high resolution ^37^. The total number of Mg^2+^ ions in non-ribosome RNA crystal structures at worse than 2.1-Å resolution is very low. The 1.7-Å structure 6YL5 of the SK209-2-6 SAM/SAH riboswitch, crystallized in a buffer containing 10 mM MgSO4 and 50 mM sodium cacodylate, resolved no Mg^2+^ but a few Na^+^ ions forming inner-shell coordination with the RNA ^21^.

A variety of computational approaches have been developed to predict metal ion binding sites in RNA structures or applied to elucidate the roles of Mg^2+^. For example, MetalionRNA (http://metalionrna.genesilico.pl/) uses a statistical potential to predict metal ions inside RNA structures ^38^; MCTBI (http://rna.physics.missouri.edu/MCTBI) predicts tightly bound ions by Monte Carlo sampling ^39, 40^; and MgNET is a convolutional neural network model trained on a set of crystal structures containing RNA and Mg^2+^ ions ^41^. Molecular dynamics (MD) simulations have been used to identify or characterize Mg^2+^ binding sites in RNA ^42–49^. When Mg^2+^ ions are initially placed in the solvent ^42, 44, 48, 49^, it can be difficult for them to form inner-shell coordination with RNA during typical MD simulation times, due to the high barrier for dehydrating Mg^2+^ ^44^; a surplus of counterions may add to the difficulty due to ion competition ^43^. Enhanced sampling can yield relative affinities of Mg^2+^ for different inner-shell sites, but requires prior knowledge of these sites ^43^. By initially placing Mg^2+^ ions at sites predicted by a structure-based method such as MCTBI, we have demonstrated success in achieving Mg^2+^-RNA inner-shell coordination in MD simulations ^46^. MD simulations have also shown that Mg^2+^ can promote the conformational transition of an RNA ^26^, quench conformational fluctuations ^47, 48^, and stabilize ligand binding ^46^.

Here we carried out 120 μs of MD simulations on the env9b SAM/SAH riboswitch (Table S1) to uncover how Mg^2+^ mediates ligand binding and conformational transition. Eleven Mg^2+^ ions stably form inner-shell coordination with backbone phosphates of the riboswitch, whether it is in the apo form or bound with SAM or SAH; nine of these sites are common among the three forms. Six of the common sites line either side of a groove that provides the entryway for the ligand. In the apo form, Mg^2+^ ions at these sites widen the groove and thus pre-organize the riboswitch for ligand binding. Once the ligand is bound, three of these Mg^2+^ ions can alternately form outer-shell coordination with the carboxy moiety of the ligands and also stabilize an additional U16-ligand hydrogen bond. These interactions have the effect of diminishing the selectivity between SAM and SAH. A unique inner-shell Mg^2+^ ion in the apo form maintains a curved shape for the G38-G39-A40 backbone, thereby loosening their basepairing with C15-C14-U12 and facilitating the release of the SD sequence for ribosome binding.

## RESULTS

### Nucleotide-ligand interaction energies and ligand exposure clarify why the riboswitch is not selective between SAM and SAH

Our MD simulations used the SAH-bound NMR structure 6HAG as the initial structure. We replaced SAH with SAM to generate the SAM-bound form and removed SAH to generate the apo form. We ran four replicate 1-μs simulations of the liganded forms starting from each of the 10 models in 6HAG, and calculated the interaction energies of the 43 nucleotides of the riboswitch with the ligand by the MM/GBSA method ^50^ (Figure 2A). Consistent with the binding pocket characterized by the NMR and crystal structures (Fig. 1C), only five nucleotides, C8, G9, U16, G17, and G38, make major contributions to the ligand binding energy. The interaction energies of each nucleotide with the two ligands are very close; however, they do favor SAM binding slightly but systematically, with the exception of U16. The general favorability of SAM can be attributed to long-range electrostatic attraction between the positive charge on its sulfur center and the RNA phosphates. The reason for the counteraction of U16 will be presented below. These interaction energy results explain why the riboswitch has only a modestly higher binding affinity for SAM than for SAH (*K_D_* = 1.5 μM and 3.7 μM for the two ligands) ^20^.

**Figure 1.**
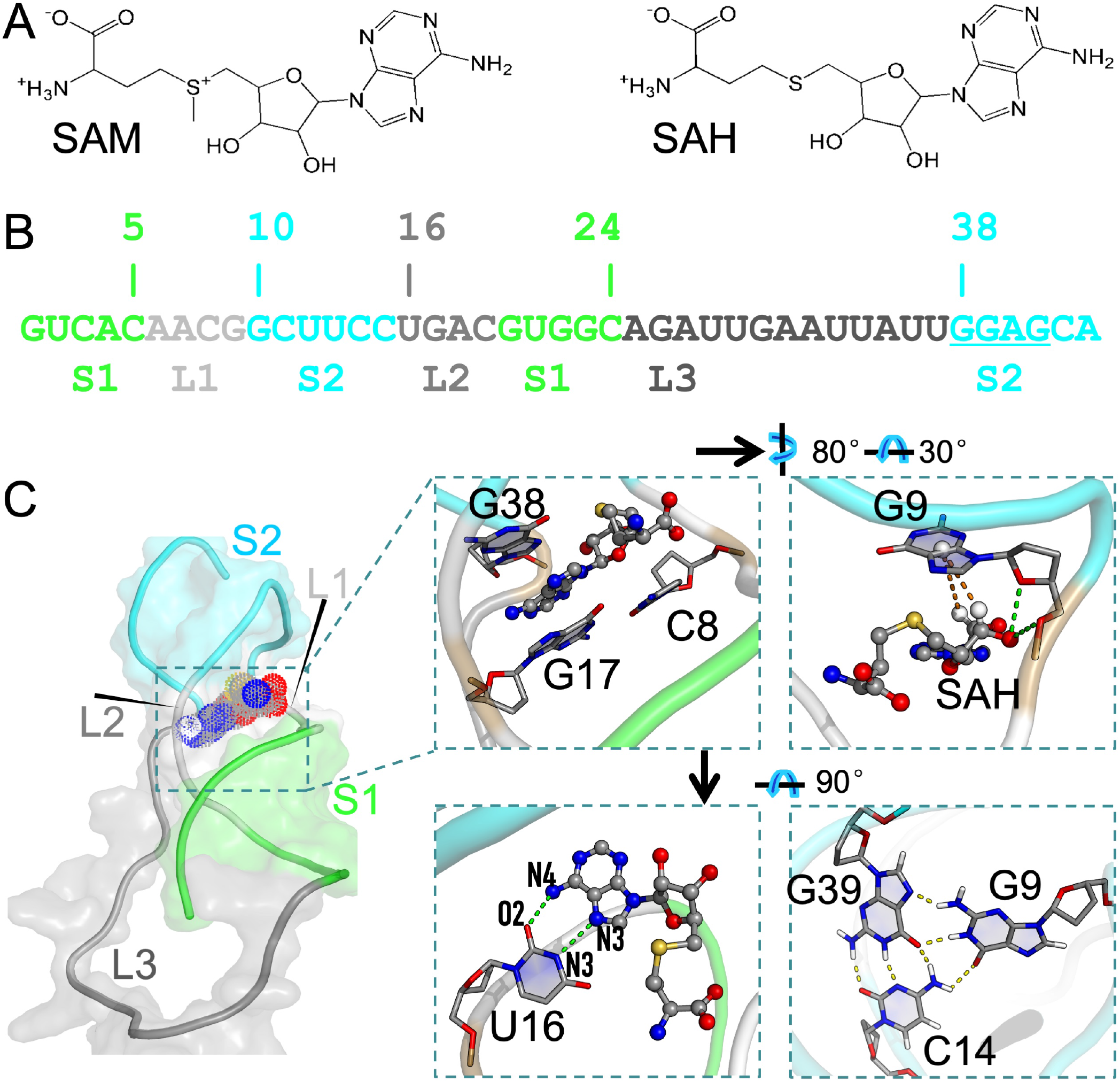
The structures of the SAM/SAH riboswitch and its cognate ligands. (A) Chemical structures of SAM (left) and SAH (right). (B) Sequence and secondary structure of the riboswitch. (C) Structure of the SAH-riboswitch complex (model 5 in 6HAG). Stacking and in-plane hydrogen bonding are highlighted in three zoomed views. The fourth zoomed view shows the base triple formed by G9, C14, and G39.

The MD simulations started from the different NMR models produced very similar interaction energies. We calculated Pearson’s correlation coefficient (*r*) between the interaction energies from any two starting models (Figure S1). For both ligands, all the pairwise correlation coefficients are close to 1, with a minimum of 0.93 between any two models. The 10 models mostly differ in the conformations of the L3 loop (A25 to U37). As can be seen in Figure 2A, except for the last two nucleotides (U36 and U37), L3 practically contributes no interaction energy with either ligand. Given the null effect of the starting model on nucleotide-ligand interaction energies, we limited to a single model in subsequent simulations. We chose model 5 because it has the smallest root-mean-square-deviation from the average structure of the 10 models.

**Figure 2.**
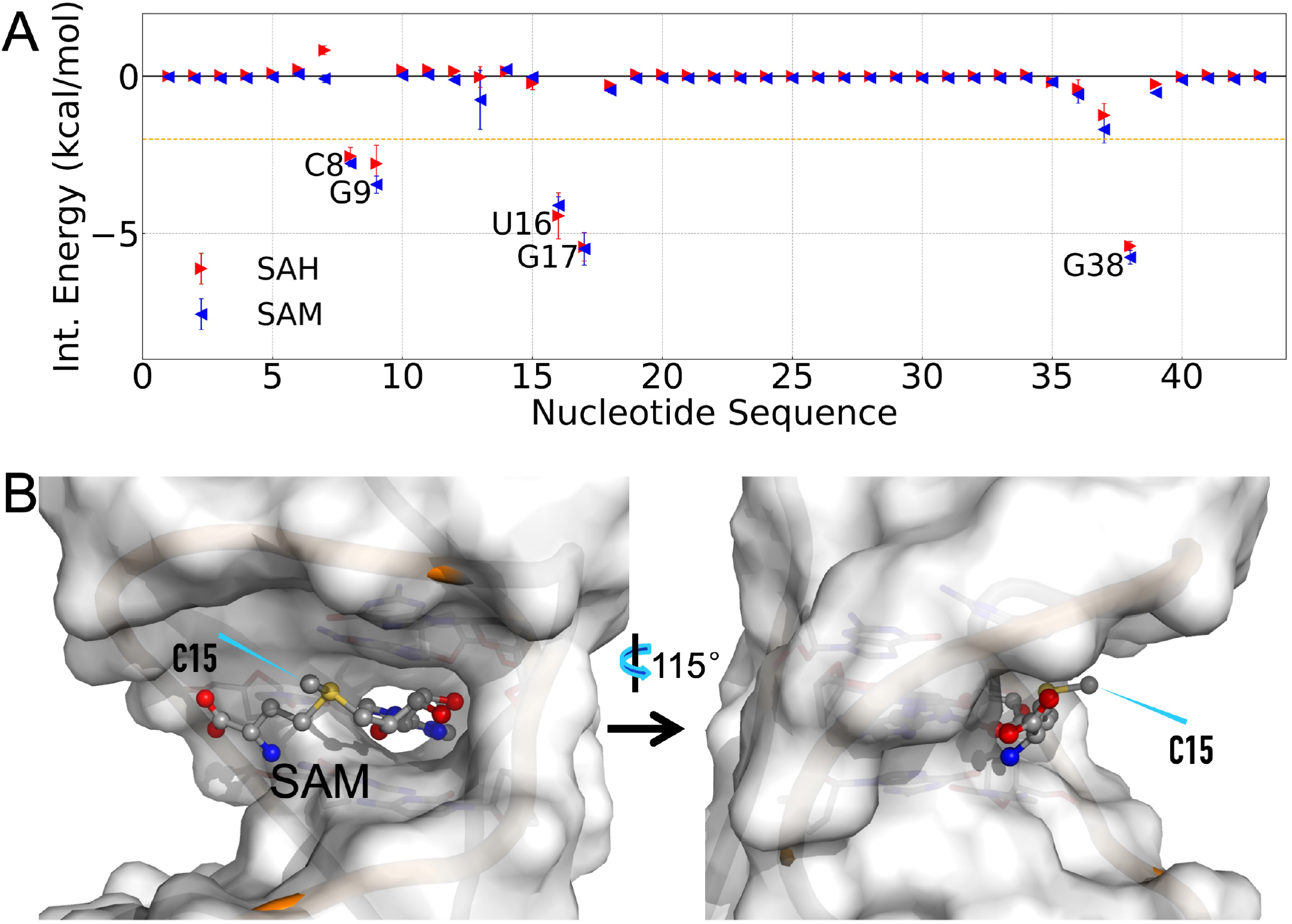
Lack of selectivity between SAM and SAH and a major reason. (A) Interaction energies of individual nucleotides with the ligands. For each nucleotide, triangle and error bar represent the mean and standard deviation, respectively, calculated among results from 10 starting models. A horizontal line at −2.0 kcal/mol separates out the five pocket-lining nucleotides. (B) The solvent exposure of the SAM aminocarboxypropyl group, especially the methyl (C15), illustrated by a snapshot from the MD simulations under saturating Mg^2+^. The backbone of the groove-lining nucleotides 5-8 and 12-16 is in orange.

While the 5’-deoxyadenosyl group of the ligand is buried in the binding pocket, the aminocarboxypropyl group and the sulfur center are exposed to a groove defined by nucleotides 5-8 and 12-16 (Figure 2B). In the MD simulations, the aminocarboxypropyl group is extremely mobile (see below); the ligand bends around the sulfur center so the aminocarboxypropyl group directs back toward the RNA, leaving the methyl on the sulfur center of SAM to be the most exposed to the solvent (Figure 2B). Therefore, although this methyl plays a central role in distinguishing from SAH in SAM-sensing riboswitches, it loses this ability in the SAM/SAH riboswitch by projecting into the solvent. The mobility of the aminocarboxypropyl group is supported by the fact that this group is oriented in different directions in 6HAG and 6YL5.

The high flexibility of Loop L3, already evident from its different conformations in the NMR models, was directly assessed by ^1^H-^13^C heteronuclear Overhauser effects (hetNOE) in the SAH-bound form ^20^ (Figure S2A). From the MD simulations, we calculated the root-mean-square-fluctuations (RMSFs) of the corresponding atoms, i.e., aliphatic H1’/C1’ or aromatic H6/C6 (for C and U nucleotides) and H8/C8 (for A and G nucleotides) (Figure S2B). The results agree well with the hetNOE data. That is, L3 and terminal nucleotides show elevated flexibilities, and both U13 and U37 show higher flexibilities than their immediate neighbors. The flexibility of U13 can be explained by the fact that, although it is a part of stem S2, it only participates in a base triple with the U12:A40 base pair. As for U37, our MD simulations reveal that this base samples two alternative poses, as further described below. Relative to the liganded forms, the apo form exhibits higher RMSFs at both U13 and U37, capturing to some extent its reported conformational heterogeneity ^11, 20, 21^.

### SAM/SAH riboswitch can harbor over 10 inner-shell Mg^2+^ ions

Following our previous study ^46^, we tested three protocols for the initial placement of Mg^2+^ ions in our RNA systems. Using MCTBI ^40^, we identified 25 putative tight-binding sites and placed Mg^2+^ ions at all these sites in the initial structure for MD simulations. Using the Leap module in AMBER18 ^51^, we added 21 Mg^2+^ ions [Leap(21)], enough to neutralize the charge on the RNA, in the solvent. The third protocol also relied on Leap but with 41 Mg^2+^ ions added [Leap(41)], thereby creating a surplus of counterions. With the MCTBI protocol, we found 11 Mg^2+^ ions stably bound to inner-shell sites in each of the three forms (apo, SAH-bound, and SAM-bound) of the riboswitch throughout the simulations. With the Leap(21) protocol, the numbers of inner-shell ions reduced to 6, 9, and 8, respectively; about one third of these ions are at the same sites as found in the simulations started with the MCTBI protocol. With the Leap(41) protocol, not a single inner-shell ion was found, due to ion competition and similar to many other MD simulation studies ^42–44, 48, 49^. Below we focus on the results from the simulations started with the MCTBI protocol, to model the condition where the RNA is saturated with Mg^2+^ and to draw the most contrast with the Mg^2+^-free condition.

To identify the RNA atoms that form inner or outer-shell coordination with Mg^2+^ ions, we calculated the radial distribution functions (RDFs) of Mg^2+^ around oxygen and nitrogen atoms on the backbone and bases. Inner and outer-shell coordination can be identified from RDF peaks at 2.0 Å and 4.3 Å, respectively (Figure 3A and Figure S3). Inner-shell coordination is nearly exclusively formed with OP1 and OP2, with OP2 favored about 2-fold over OP1. Second-shell coordination is most frequently formed with OP1, OP2, O3’, and O5’ on the backbone, and less frequently with bases. Of the latter, the N7 atom of the A base, N7 and O6 of G, and O4 of U are the most frequent. These statistics obtained from our MD simulations of the riboswitch in the apo and liganded forms are in remarkable agreement with Mg^2+^ coordination frequencies tabulated from crystal structures by Zheng et al. ^37^.

**Figure 3.**
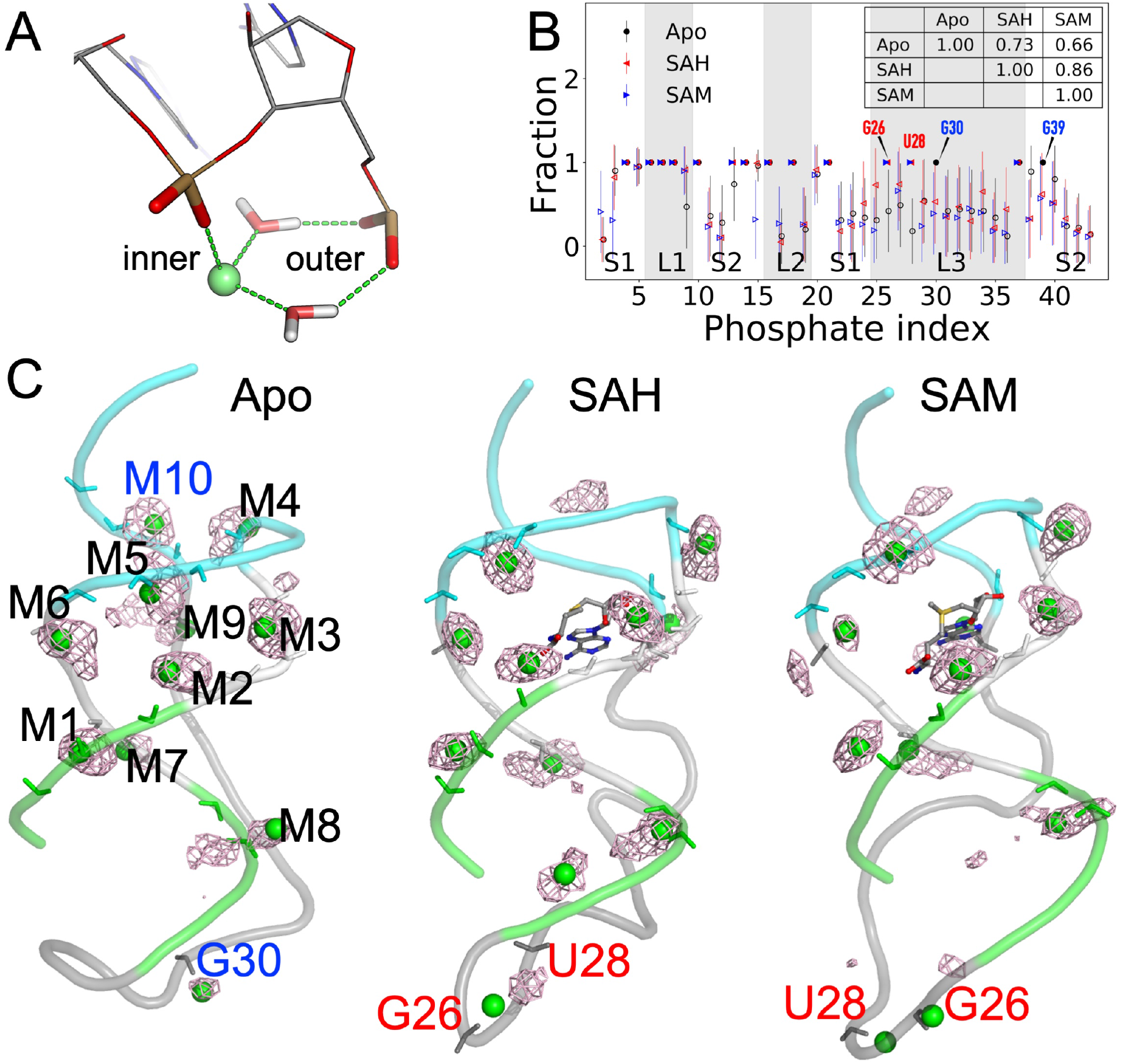
Inner-shell Mg^2+^ ions in MD simulations of the apo and two liganded forms. (A) Illustration of a Mg^2+^ ion forming both inner-shell coordination with one phosphate and outer-shell coordination with an adjacent phosphate. Hydrogen bonds are indicated by dashed lines. (B) The fraction of frames where a phosphate forms inner or outer-shell coordination. Inner-shell coordination, once formed, is stable in the MD simulations (fraction = 1; solid symbols). For outer-shell fractions, open symbols and error bars represent the means and standard deviations, respectively, calculated among results from four replicate simulations. Inset table: correlation coefficients between any two forms of the riboswitch. (C) Densities of Mg^2+^ ions, displayed as mesh and superimposed on a representative snapshot from the MD simulations. The inner-shell Mg^2+^ ions are shown as green spheres, and the coordinating phosphates are shown as sticks.

In Figure 3B, we present the fraction of MD frames where each nucleotide forms inner or outer-shell coordination. Most of the inner-shell nucleotides are found in the first 21 nucleotides, containing stem S1, loops L1 and L2, and the 5’ strand of stem S2, and are largely conserved among the apo and two liganded forms. The coordination patterns in the remaining 22 nucleotides show differences between the apo form and the two liganded forms. Overall, the correlations of the fraction values are strong between the two liganded forms (*r* = 0.86) but moderately reduced between either liganded form and apo (inset table). In Figure 3C, we display the densities of Mg^2+^ ions around the RNA, superimposed on a representative snapshot of inner-shell Mg^2+^ ions.

For each inner-shell Mg^2+^ ion, we identified its coordinating phosphates by calculating the distributions of their distances (Figures S4 and S5). We name Mg^2+^ ions that form inner-shell coordination outside the flexible portion of loop L3 as M1 to M10 (Figure 3C). A few Mg^2+^ ions form inner-shell coordination simultaneously with two adjacent phosphates, in a bidentate configuration ^23^, such as M3 with A7 and C8, or only with a single phosphate, such as M7 with A18. However, most inner-shell Mg^2+^ ions form outer-shell coordination with an adjacent phosphate. Outer-shell coordination occurs most frequently with both OP1 and OP2 of the adjacent phosphate via two bridging water molecules, as illustrated in Figure 3A, but can also occur with only OP1 or OP2, via either one or two bridging water molecules. M1-M8 coordinate with the first 21 nucleotides and are largely conserved among the simulations of the three forms of the riboswitch (Figure S4). The flexible portion of L3 harbors one inner-shell Mg^2+^ ion, coordinating with G30, in the apo form, but two other inner-shell Mg^2+^ ions, coordinating either G26 or U28, in the liganded forms (Figure S5). These Mg^2+^ ions move with the highly flexible L3, and therefore their densities are smeared out in space. U37 (nominally on L3) harbors the last conserved inner-shell Mg^2+^ ion M9, and the 3’ strand of stem S2 harbors one last inner-shell Mg^2+^ ion, M10, only in the apo form. Below we present further details of these Mg^2+^ ions and their structural and functional consequences.

### Inner-shell Mg^2+^ ions widen a groove and pre-organize the riboswitch for ligand entry

As presented above, the 5’-deoxyadenosyl group of the ligand is buried in the binding pocket, with the groove defined by nucleotides 5-8 and 12-16 providing the entryway. We used the distance, *d*_P6-P14_, between the phosphorus atoms of A6 and C14, to measure the groove width (Figure 4A). The groove width increases upon ligand binding, both without or with saturating Mg^2+^ (Figure 4B). Interestingly, in the apo form, the mean groove width is increased, from 10.9 Å without Mg^2+^ to 12.7 Å under saturating Mg^2+^. The resulting groove width is comparable to those in the liganded forms without Mg^2+^ (mean *d*_P6-P14_ at 12.9 Å and 12.1 Å, respectively, for the SAH and SAM-bound forms). Mg^2+^ saturation also results in groove widening in the liganded forms.

**Figure 4.**
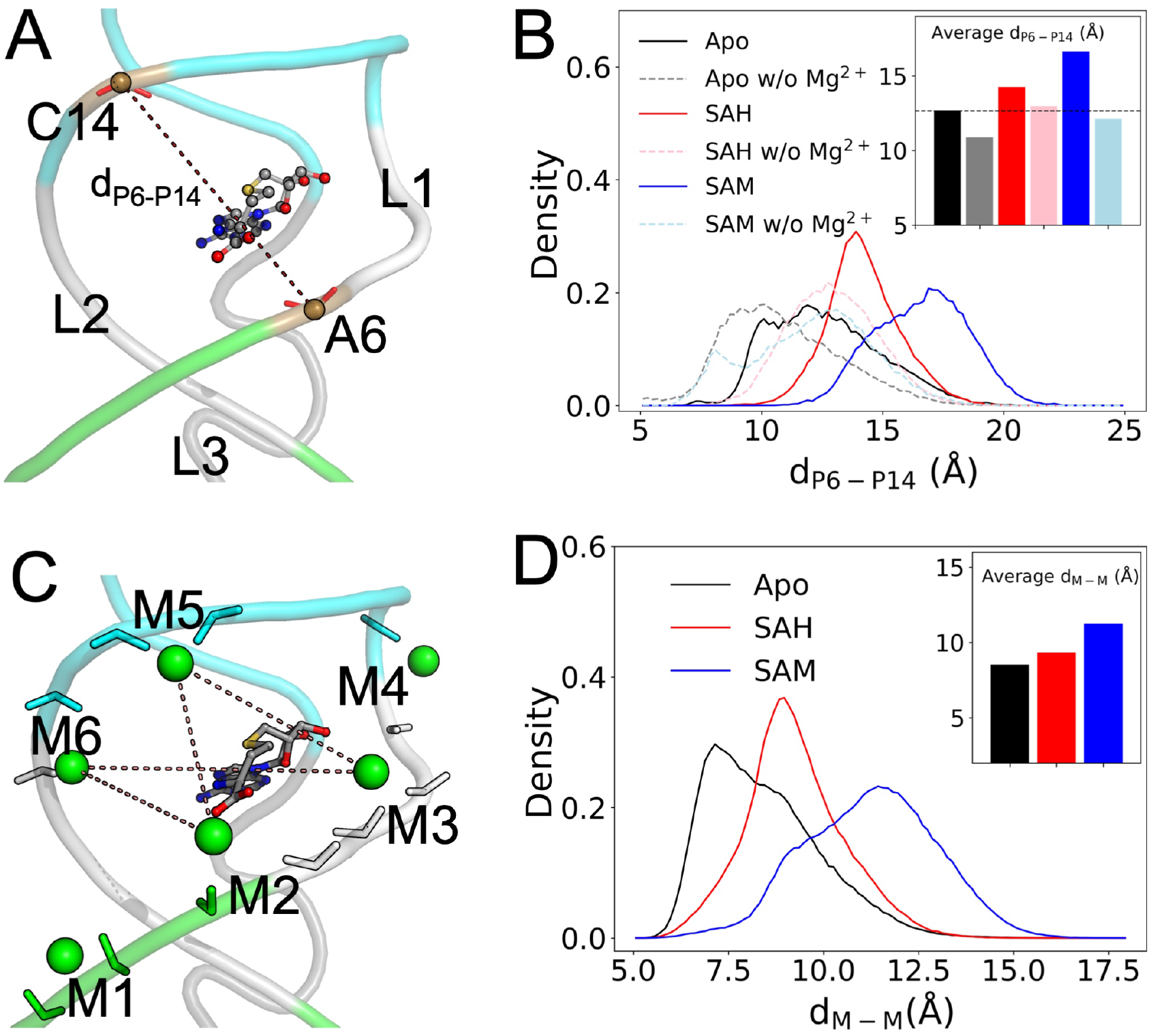
Groove widening upon ligand binding and by inner-shell Mg^2+^ ions. (A) The distance, *d*_P6-P14_, for measuring the groove width. (B) Distributions of *d*_P6-P14_ in the simulations of the three forms of the riboswitch without (labeled as “w/o”) or with saturating Mg^2+^. Inset: average values for six systems. (C) Six inner-shell Mg^2+^ ions lining the two sides of the groove. (D) Distributions of the minimum distance between M2 and M3 on one side of the groove and M5 and M6 on the opposite side. Inset: average values for the three systems.

It thus appears that inner-shell Mg^2+^ ions widen the ligand-entry groove and thereby pre-organize the riboswitch for ligand entry. Six conserved inner-shell Mg^2+^ ions, M1-M6, line the two sides of this groove (Figure 4C). The minimum distance between M2 and M3 (coordinating with C5-C8) on one side and M5 and M6 (coordinating with U13-U16) on the opposite side mirrors the groove width (Figure 4B, D). Likely the electrostatic repulsion between M1-M3 and M4-M6 on the opposite sides of the groove contributes to the groove widening upon Mg^2+^ saturation.

We can even further speculate that, in the apo form, inner-shell Mg^2+^ ions M1-M6 may hold the incoming ligand at the groove by electrostatic attraction with its carboxy moiety. The 5’-deoxyadenosyl group would then explore the groove and get into the binding pocket via the pre-widened entryway.

### Inner-shell Mg^2+^ ions form outer-shell coordination with ligands and stabilize U16-ligand hydrogen bonding

The aminocarboxypropyl group of the ligands is exposed to the ligand-entry groove and is extremely mobile in the MD simulations. The carboxy moiety can potentially form outer-shell coordination with three of the groove-lining Mg^2+^ ions: M2, M5, and M6 (Figures 4C and 5A). We calculated the shorter of the distances from the two oxygen atoms on the ligand carboxy to each of these three Mg^2+^ ions (Figure S6A, B). There are indeed substantial fractions of MD frames where the ligand carboxy is within the 2.5 to 5 Å range for outer-shell coordination with M6, M5, and M2 (Figure 5B). For the SAH-bound form, the outer-shell coordination fractions with these Mg^2+^ ions are 77.7%, 34.7%, and 13.6%, respectively. Coordination with M6 and M5 can occur simultaneously, but coordination with either M6 or M5 and coordination with M2 are mutually exclusive, as they are located on opposite sides of the groove. The high propensity that the ligand carboxy moiety forms outer-shell coordination with at least one of the groove-lining Mg^2+^ ions buttresses the foregoing speculation about their potential role in holding the ligand at the groove prior to binding.

**Figure 5.**
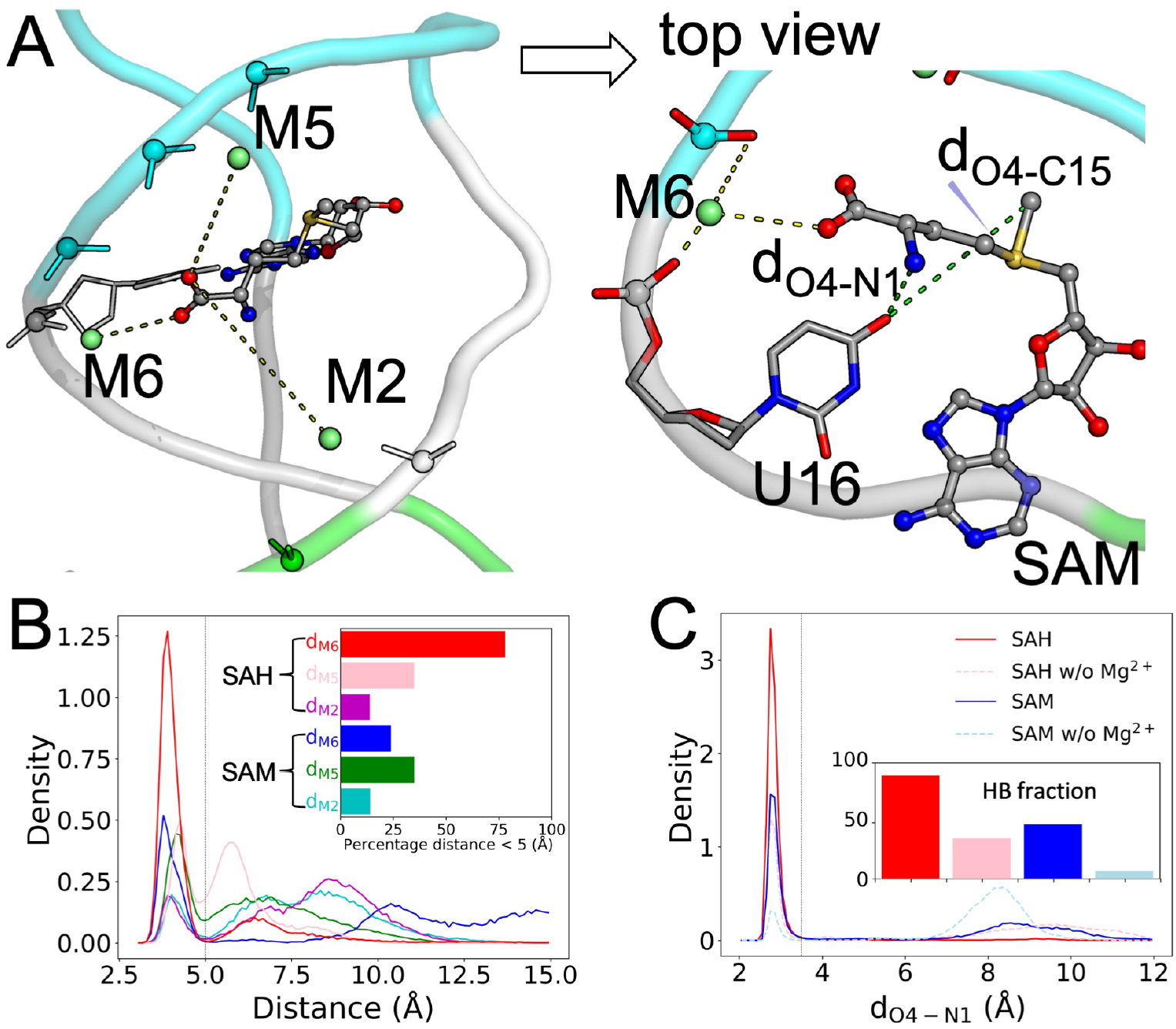
Outer-shell coordination with ligands and stabilization of U16-ligand hydrogen bond by M2, M5, and M6. (A) The positions of M2, M5, and M6 relative to the ligand carboxy moiety. Top view: M6 forms inner-shell coordination with C15 and outer-shell coordination with both U16 and SAM carboxy. Also shown are the distances between the U16 O4 atom and SAM N1 and C15 atoms. (B) Distributions of the distances between the ligand carboxy and M2, M5, and M6. A vertical line at 5 Å indicates the cutoff for outer-shell coordination. Inset: fractions of frames forming outer-shell coordination. (C) Distributions of *d*_O4-N1_. A vertical line at 3.5 Å indicates the cutoff for hydrogen bond formation. Inset: fractions of frames forming a U16-ligand N1 hydrogen bond.

For the SAM-bound form, the coordination fraction with M6 is significantly reduced, to 23.7%, while those with M5 and M2 are similar to the SAH counterparts. Whereas the SAH-bound form has similarly high propensities for outer-shell coordination with M6 in all the four replicate simulations, the SAM-bound form does so in only one (MD4) of the four replicate simulations. Therefore the inner-shell Mg^2+^ ions stabilize ligand binding by forming outer-shell coordination with the ligand carboxy. However, the stabilizing effect, specifically from M6, is greater for SAH than for SAM, thereby countering the general favorability of SAM (Figure 2A) and diminishing potential selectivity against SAH.

We have noted above that U16 also makes a greater contribution to the binding energy for SAH than to the counterpart for SAM. The counteractions of M6 and U16 are closely linked, as we now explain. In the static NMR structure 6HAG, the O4 atom of U16 is near but outside the hydrogen-bonding distance (3.5 Å) from the amino N1 atom of the ligand ^20^. However, in the MD simulations, *d*_O4-N1_ (Figure 5A, top view) frequently comes into the hydrogen-bonding range (Figure S6C, D). The fractions of MD frames forming the U16-ligand amino hydrogen bond are 88.8% for SAH and 48.1% for SAM (Figure 5C). The higher hydrogen-bonding fraction for SAH accounts for the greater contribution of U16 to the binding energy for this ligand.

For SAM, the methyl (C15 atom) on the sulfur center can potentially clash with the U16 O4 atom (Figure 5A, top view). Indeed, *d*_O4-N1_ and *d*_O4-C15_ are anticorrelated: when SAM C15 gets close to U16 O4, SAM N1 moves away, and vice versa (Figure S6E; *r* = −0.51). Therefore the methyl can interfere with U16-ligand amino hydrogen bonding, thereby explaining the much higher propensity of forming this hydrogen bond by SAH and the greater contribution of U16 to the binding energy of this ligand.

When this hydrogen bond with U16 is formed, it places the carboxy moiety in a position to form outer-shell coordination with M6. Therefore, when the methyl of SAM interferes with U16-ligand amide hydrogen bonding, it simultaneously interferes with M6-ligand carboxy outer-shell coordination. In simulation MD4 where SAM forms M6-ligand carboxy coordination with 94.9% probability (Figure S6B), it also forms the U16-ligand amide hydrogen bond with 98.6% probability. In contrast, in simulations MD1-MD3 where coordination with M6 is never formed, the probability for the hydrogen bond drops to 31.3% (Figure S6D). Moreover, while the M6-ligand carboxy distance in these three simulations never reaches the outer-shell coordination cutoff, it shows a strong correlation with the U16-ligand amide distance (Figure S6B, D; *r* = 0.87).

In short, the interactions of the ligands and Mg^2+^ ion M6 with U16 are highly cooperative, with a strong tendency to form or break at the same time. As a last testament to this cooperativity, without Mg^2+^, the probabilities for forming the U16-ligand amino hydrogen bond are dramatically reduced for both SAH and SAM (Figure 5C and Figure S6D).

### An inner-shell Mg^2+^ ion in the apo form facilitates the release of the SD sequence

Upon ligand binding, the U37 nucleobase is extruded from the ligand-binding pocket. In the MD simulations, this base adopts two alternative poses (Figure 6A, B). We monitored the movement of this nucleobase by calculating its distance (*d*_pk-B37_) from the center of the binding pocket (defined by the pocket-lining bases C8, G9, U16, G17, and G38) (Figure 6C). The distribution of *d*_pk-B37_ exhibits two peaks, corresponding to the two alternative poses, for either the SAH or SAM-bound form. These alternative poses explain why its RMSF is higher than those of its immediate neighbors (Figure S2B). The upstream neighbor, U36, tends to form a hydrogen bond, via its 2’-OH, with either the N4 or N5 atom of the ligand base (Figure 6D, zoomed view on the right). The downstream neighbor, G38, base pairs with C15 as part of stem S2.

**Figure 6.**
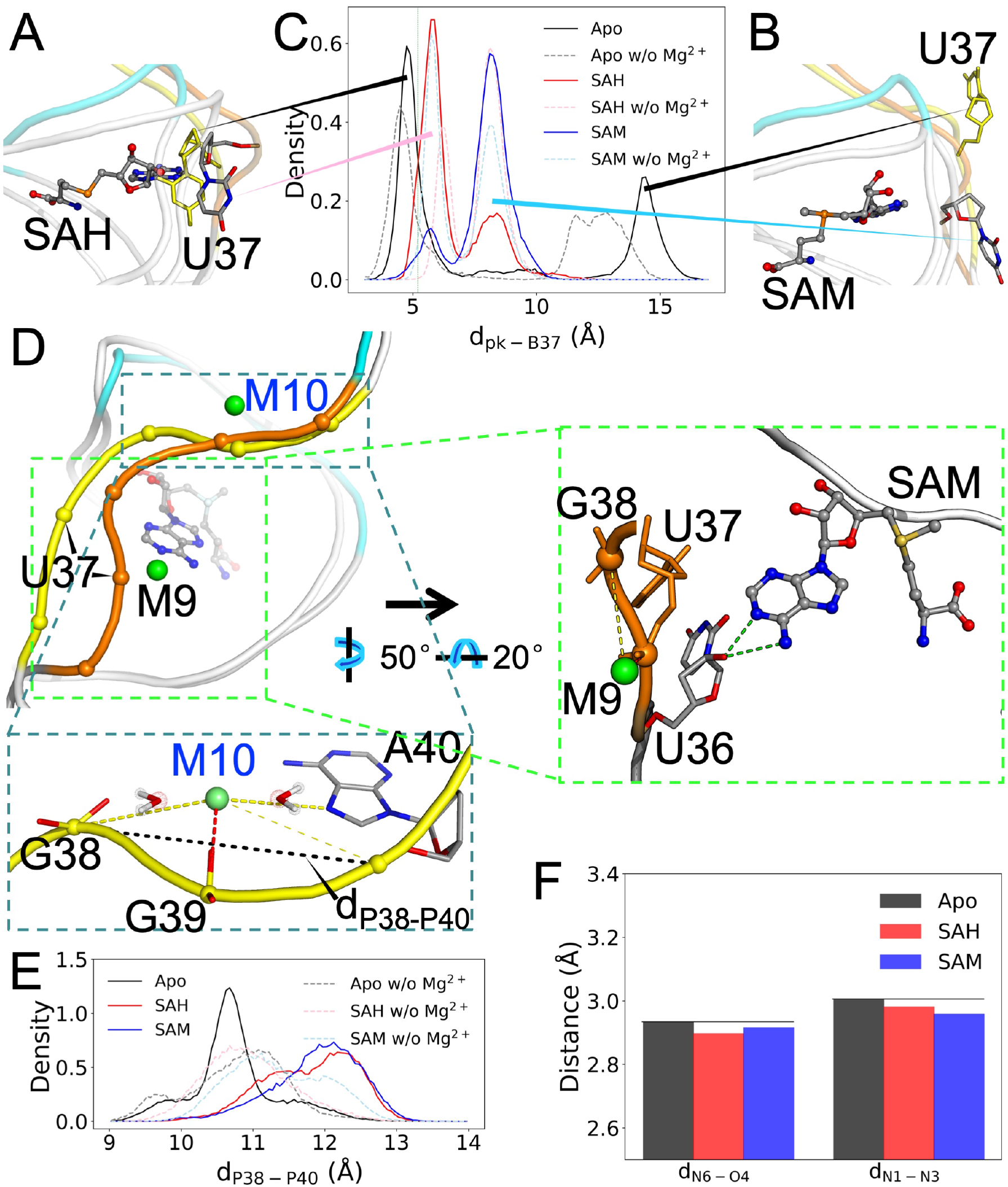
M10 facilitates the release of the SD sequence in the apo form. (A) The U37 nucleobase is extruded upon ligand binding (C, N, and O atoms in gray, blue, and red, respectively), but can move into the ligand-binding pocket in the apo form (all yellow). (B) An alternative pose for U37, just outside the binding pocket in the liganded form but farther out in the apo form. (C) Distributions of *d*_pk-B37_, the distance from the U37 nucleobase to the center of the binding pocket. (D) Backbone shape of nucleotides 36-40, in orange for the SAM-bound form and yellow for the apo form. Zoomed view of the SAM-bound form on the right: outer-shell coordination of M9 with G38 and hydrogen bonding between U36 2’-OH and SAM N4 or N5. Zoomed view of the apo form at the bottom: M10 forms inner-shell coordination with G39 and outer-shell coordination with both G38 and A40. Also shown is the distance between the phosphates of G38 and A40. (E) Distributions of *d*_P38-P40_. (F) Average distances between hydrogen-bonding donors and acceptors on the base-pair partners A40 and U12.

In the simulations of the apo form, the U37 nucleobase moves either into the binding pocket (Figure 6A; *d*_pk-B37_ ~4.7 Å) or far away from the binding pocket (Figure 6B; *d*_pk-B37_ ~14.4 Å), spending roughly equal time in the two positions (Figure 6C). The large distance between these two positions of U37 contributes to the conformational heterogeneity of the apo form.

The ligand-forced extrusion of U37 and the stacking of G38 against the ligand base (Figure 1C) result in changes in both the backbone curvature and the Mg^2+^ coordination pattern on the downstream side. Relative to the apo form, the phosphates of U37 and G38 are brought closer, and the backbone of nucleotides G38-G39-A40 is straightened (Figure 6D and zoomed view at the bottom). We measured the latter effect by calculating the distance, *d*_P38-P40_, between the phosphorus atoms of G38 and A40 (Figure 6E). The peak distance increases from 10.6 Å in the apo form to 12.4 Å in both of the liganded forms.

Mg^2+^ ion M9 forms inner-shell coordination with the U37 phosphate in all the three forms of the riboswitch (Figure S5 and Figure 6D). M9 also forms outer-shell coordination with G38 part of the time in the liganded forms due to the closer U37-G38 distance (Figure S5, Figure 3B, and Figure 6D, zoomed view on the right), but not all in the apo form. Instead, the curved backbone of apo G38-G39-A40 creates an inner-shell Mg^2+^ site that is unique to the apo form. M10 forms inner-shell coordination with G39 and outer-shell coordination with both G38 and A40; for A40, outer-shell coordination can occur via either the phosphate or the base N7 atom (Figure S5 and Figure 6D, zoomed view at the bottom). Note that the apo G38-G39-A40 backbone is curved and M10 is bound all the time, regardless of whether U37 takes its position inside or away from the binding pocket; the distribution of *d*_P38-P40_ has only a single peak. Therefore the curved shape may be intrinsic to G38-G39-A40, stabilized by the inner-shell Mg^2+^ ion M10. Without Mg^2+^, the *d*_P38-P40_ distribution in the apo form no longer shows distinction from those in the liganded forms (Figure 6E).

Because the M10 site exists exclusively in the apo form and resides completely inside the SD sequence, we suspected that it might play a direct functional role. One possibility is that M10 maintains G38-G39-A40 in an autonomous mode such that their base-pairing with the upstream nucleotides C15-C14-U12 is weakened. Figure 6F shows that, indeed, for the base pair between A40 and U12, the donor-acceptor distances are longer in the apo form. Therefore M10 can directly facilitate the release of the DS sequence to initiate translation.

## DISCUSSION

We have carried out extensive MD simulations to investigate the essential roles of Mg^2+^ in the ligand binding and conformational transition of the SAM/SAH riboswitch. We identified 11 inner-shell Mg^2+^ ions each in the apo form and the SAM and SAH-bound forms. Six of the common Mg^2+^ ions (M1 to M6) line the ligand-entry groove to widen it, thereby pre-organizing the riboswitch for ligand binding. M2, M5, and M6 alternately form outer-shell coordination with the ligands. In addition, M6 stabilizes the U16-ligand amide hydrogen bond. These interactions occur with reduced probability for SAM due to the interference of its methyl, thereby countering the general favorability of this ligand over SAH and diminishing the selectivity between these two ligands. One Mg^2+^ ion, M10, unique to the apo form maintains the SD sequence in a curved conformation and weakens its base-pairing with upstream nucleotides, thereby facilitating its release for ribosome binding.

Key aspects of our MD results are validated by experimental observations. For example, the flexibility profiles calculated from the MD simulations agree well with ^1^H-^13^C hetNOE data ^20^. The extreme mobility of the ligand carboxy moiety seen in our MD simulations is supported by its different orientations in the NMR structure 6HAG and crystal structure 6YL5 ^20, 21^ (Figure S7A). The probabilities of different RNA atoms forming inner and outer-shell coordination with Mg^2+^ match those tabulated from crystal structures ^37^. As further validation, we compared the three inner-shell Na^+^ ions in 6YL5 with our inner-shell Mg^2+^ sites. One Na^+^ ion forms inner-shell coordination with the ligand carboxy and outer-shell coordination with A6 and A7 (env9b riboswitch numbering); this Na^+^ is similar to our M2. The second Na^+^ ion forms inner-shell coordination with G20 and outer-shell coordination with C19, close to our M8. The third Na^+^ ion forms inner-shell coordination with G39 and outer-shell coordination with A40 (via both phosphate and N7). Interestingly, this Na^+^ ion is very much like our M10, except that ours is found in the apo form. Comparing backbone shapes of the nearby nucleotides, we find that 6LY5 is more curved than 6HAG (Figure S7A), to a similar extent as our apo form (Figure S7B). It looks as if the removal of the flexible L3 in the crystal structure of the SAH-bound form reduces restraints on G39 and A40 so they behave as if in the apo form.

Our MD simulations have generated unique mechanistic insights. In particular, the solvent exposure of the methyl on SAM leads to similar interaction energies for SAM and SAH with the riboswitch nucleotides, yet the positive charge on the sulfur center still generally favors SAM over SAH. It is Mg^2+^ ions that provide compensatory effects for SAH and dimmish the selectivity between the two ligands. Without the methyl, M6 is better able to form outer-shell coordination with the SAH carboxy and stabilize the U16-SAH amide hydrogen bond. We also show that Mg^2+^ ions widen the ligand-entry groove so to reduce the energy barrier for entering the ligand-binding pocket. We further speculate that these Mg^2+^ ions can potentially hold the ligand, via outer-shell coordination with its carboxy moiety, to give the ligand more chance to explore the groove and enter the ligand-binding pocket. Lastly, while Mg^2+^ ions are generally known to stabilize RNA structures including helical elements, our characterization of M10 in the apo form shows that they can also specifically interact with one strand of a helical element and peel it away from the complementary strand. The end result, for the SAM/SAH riboswitch, is the release of the SD sequence.

Despite their essential roles illustrated here, Mg^2+^ is difficult to identify and can thus be dubbed the “dark” metal ion in RNA research. The present study, along with our previous work ^46^, has demonstrated that placing Mg^2+^ ions initially according to a structure-based prediction method such as MCTBI ^40^ is an effective protocol for producing inner-shell ions in conventional MD simulations. We hope that this protocol and further developments will make MD simulations an even more powerful technique for characterizing both the structural determinants and the functional consequences of Mg^2+^ coordination.

## COMPUTATIONAL METHODS

### Preparation of RNA systems

The initial SAH-bound structure of the SAM/SAH riboswitch was from the NMR structure 6HAG ^20^. The SAH ligand was replaced by SAM to generate the SAM-bound structure and removed to generate the apo structure. The original hydrogen atoms on the RNA molecule were removed and re-added by using the Leap module in AMBER18 ^51^. For each of the three forms (SAH or SAM-bound or apo) of the riboswitch, four types of Mg^2+^ initial placement were applied (Table S1): (i) without any Mg^2+^; (ii) addition of 25 Mg^2+^ ions at sites predicted by MCTBI ^40^; (iii) addition of 21 Mg^2+^ ions (enough to neutralize the RNA molecule) as part of the solvation step using Leap; and (iv) addition of 41 Mg^2+^ ions using Leap. Type (iv) was applied to all the 10 models in 6HAG for the two liganded forms; in all other cases only model 5 was prepared. In the solvation step, the RNA [plus Mg^2+^ in the case of type (ii)] was placed in a truncated octahedron periodic box of TIP3P ^52^ water molecules. Mg^2+^ [in the cases of types (iii) and (iv)], neutralizing Na^+^ or Cl^-^ ions, and 0.15 M NaCl [except type (i)] were also added in the solvent. The distance from the RNA molecule to the edge of the box was at least 12 Å.

The force field for RNA was an improved version of AMBER ff99 ^53^, with correction for α/γ dihedrals (bsc0) ^54^ and correction for χ dihedrals (χ_OL3_) ^55^. The parameters for Mg^2+^ were from Li et al. ^56^; those for Na^+^ and Cl^-^ were from Joung and Cheatham ^57^. To generate force-field parameters for the ligands, the structures of the ligands were optimized using the Gaussian16 program ^58^ at the HF/6-31G* level. The atomic partial charges were assigned using the restrained electrostatic potential (RESP) method ^59^; other parameters were taken from the general Amber force field ^60^.

### Molecular dynamics simulations

Energy minimization and MD simulations were carried out using the AMBER18 package ^51^. To start, each system was minimized by the steepest-descent and conjugate-gradient methods, each for 2500 steps. The preparatory stage of the simulation consisted of 50 ps of temperature ramping from 100 K to 300 K at constant volume, 50 ps at 300 K and constant volume, and 50 ps at 300 K and 1.0 atm pressure, while restraining the RNA and ligand atoms with a force constants of 5 kcal/(mol·Å^2^). The equilibration stage was 1 ns at constant temperature and pressure (without restraints). The production run was carried out in four replicates for 1 μs at constant temperature and pressure. The temperature (300 K) was regulated by the Langevin thermostat ^61^, and pressure (1.0 atm) was regulated by the Berendsen barostat ^62^. Long-range electrostatic interactions were treated by the particle mesh Ewald method ^63^ with a direct-space cutoff of 12 Å, and bonds involving hydrogen atoms were constrained by the SHAKE algorithm ^64^. The time step was 2 fs. Frames were saved at 100 ps intervals for later analysis.

### Interaction energies between individual nucleotides and the ligand

The interaction energies were calculated by the MM/GBSA method ^50^ using AMBER18. Results were obtained for 5000 frames in the second 500 ns of each replicate simulation, and then averaged over four replicate simulations (or further over 10 starting models).

### Other analyses

All other analyses were done on 10000 frames in the entire 1000 ns of each replicate simulation and then averaged over four replicate simulations. RMSFs were calculated by first aligning using backbone atoms (P, O3’, O5’, C3’, C4’, C5’, excluding L3) of the riboswitch to obtain an average structure and then finding the deviations of a specific set of atoms from the average structure. The set of atoms was: (i) H1’ and C1’; or (ii) H6 and C6 (C and U nucleotides) or H8 and C8 (A and G nucleotides). RDFs were calculated from the number of Mg^2+^ ions within a distance bin (0.05-Å width) from a given RNA atom, normalized by the expected number of Mg^2+^ ions in that bin assuming uniform density. RMSFs, RDFs, distances, and hydrogen bond formation were calculated by the CPPTRAJ program ^65^. Hydrogen bonding criteria were: donor-acceptor distance < 3.5 Å and donor-H-acceptor angle > 120°.

The fraction of frames where a nucleotide makes either inner or outer-shell coordination with Mg^2+^ions was calculated using a Tcl script in VMD ^66^, based on distances between the phosphate OP1 or OP2 atom and any Mg^2+^ion, with cutoffs at 2.5 and 5 Å, respectively. The 2.5-Å cutoff was chosen because it falls well into the gap between the first and second peaks of the RDFs around OP1 and OP2 (Figure S3); The 5-Å cutoff was chosen because it is where the second peak of the RDFs first crosses the baseline. The densities of Mg^2+^ were determined using a python script importing the MDAnalysis package ^67^.

## Acknowledgments

This work was partially supported by funding from the Natural Science Foundation of Shandong Province (ZR2019MA040 to HG), the National Natural Science Foundation of China (32171249 to HG), and the US National Institutes of Health (GM118091 to HXZ).

## Author contributions

G.H. and H.X.Z designed research, conducted research, analyzed data, and wrote manuscript.

## Competing interests

The authors declare no competing interests.

## Notes

### Competing Interest Statement

The authors have declared no competing interest.

## REFERENCES

1. Pavlova, N., Kaloudas, D., Penchovsky, R. Riboswitch distribution, structure, and function in bacteria. Gene 708, 38–48 (2019).

2. Serganov, A., Nudler, E. A decade of riboswitches. Cell 152, 17–24 (2013).

3. Ray, S., Chauvier, A., Walter, N. G. Kinetics coming into focus: single-molecule microscopy of riboswitch dynamics. RNA Biol 16, 1077–1085 (2019).

4. Widom, J. R., et al. Ligand Modulates Cross-Coupling between Riboswitch Folding and Transcriptional Pausing. Mol Cell 72, 541–552 e546 (2018).

5. Bedard, A. V., Hien, E. D. M., Lafontaine, D. A. Riboswitch regulation mechanisms: RNA, metabolites and regulatory proteins. Biochim Biophys Acta Gene Regul Mech 1863, 194501 (2020).

6. Struck, A. W., Thompson, M. L., Wong, L. S., Micklefield, J. S-adenosyl-methionine-dependent methyltransferases: highly versatile enzymes in biocatalysis, biosynthesis and other biotechnological applications. Chembiochem 13, 2642–2655 (2012).

7. Ji, W., et al. Sulfonium-Based Homolytic Substitution Observed for the Radical SAM Enzyme HemN. Angew Chem Int Ed Engl 59, 8880–8884 (2020).

8. Berger, S. L. Molecular biology. The histone modification circus. Science 292, 64–65 (2001).

9. Jones, P. A., Takai, D. The role of DNA methylation in mammalian epigenetics. Science 293, 1068–1070 (2001).

10. Batey, R. T. Recognition of S-adenosylmethionine by riboswitches. Wiley Interdiscip Rev RNA 2, 299–311 (2011).

11. Weinberg, Z., et al. Comparative genomics reveals 104 candidate structured RNAs from bacteria, archaea, and their metagenomes. Genome Biol 11, R31 (2010).

12. Winkler, W. C., Nahvi, A., Sudarsan, N., Barrick, J. E., Breaker, R. R. An mRNA structure that controls gene expression by binding S-adenosylmethionine. Nat Struct Biol 10, 701–707 (2003).

13. McDaniel, B. A., Grundy, F. J., Artsimovitch, I., Henkin, T. M. Transcription termination control of the S box system: direct measurement of S-adenosylmethionine by the leader RNA. Proc Natl Acad Sci U S A 100, 3083–3088 (2003).

14. Corbino, K. A., et al. Evidence for a second class of S-adenosylmethionine riboswitches and other regulatory RNA motifs in alpha-proteobacteria. Genome Biol 6, R70 (2005).

15. Wang, J. X., Breaker, R. R. Riboswitches that sense S-adenosylmethionine and S-adenosylhomocysteine. Biochem Cell Biol 86, 157–168 (2008).

16. Weinberg, Z., et al. The aptamer core of SAM-IV riboswitches mimics the ligand-binding site of SAM-I riboswitches. RNA 14, 822–828 (2008).

17. Poiata, E., Meyer, M. M., Ames, T. D., Breaker, R. R. A variant riboswitch aptamer class for S-adenosylmethionine common in marine bacteria. RNA 15, 2046–2056 (2009).

18. Mirihana Arachchilage, G., Sherlock, M. E., Weinberg, Z., Breaker, R. R. SAM-VI RNAs selectively bind S-adenosylmethionine and exhibit similarities to SAM-III riboswitches. RNA Biol 15, 371–378 (2018).

19. Sun, A., et al. SAM-VI riboswitch structure and signature for ligand discrimination. Nat Commun 10, 5728 (2019).

20. Weickhmann, A. K., et al. The structure of the SAM/SAH-binding riboswitch. Nucleic Acids Res 47, 2654–2665 (2019).

21. Huang, L., Liao, T. W., Wang, J., Ha, T., Lilley, D. M. J. Crystal structure and ligand-induced folding of the SAM/SAH riboswitch. Nucleic Acids Res 48, 7545–7556 (2020).

22. Draper, D. E. A guide to ions and RNA structure. RNA 10, 335–343 (2004).

23. Petrov, A. S., Bowman, J. C., Harvey, S. C., Williams, L. D. Bidentate RNA-magnesium clamps: on the origin of the special role of magnesium in RNA folding. RNA 17, 291–297 (2011).

24. Batey, R. T., Williamson, J. R. Effects of polyvalent cations on the folding of an rRNA three-way junction and binding of ribosomal protein S15. RNA 4, 984–997 (1998).

25. Rangan, P., Woodson, S. A. Structural requirement for Mg2+ binding in the group I intron core. J Mol Biol 329, 229–238 (2003).

26. Bergonzo, C., Hall, K. B., Cheatham, T. E., 3rd. Divalent Ion Dependent Conformational Changes in an RNA Stem-Loop Observed by Molecular Dynamics. J Chem Theory Comput 12, 3382–3389 (2016).

27. Martinez-Monge, A., Pastor, I., Bustamante, C., Manosas, M., Ritort, F. Measurement of the specific and non-specific binding energies of Mg(2+) to RNA. Biophys J 121, 3010–3022 (2022).

28. Noeske, J., Schwalbe, H., Wohnert, J. Metal-ion binding and metal-ion induced folding of the adenine-sensing riboswitch aptamer domain. Nucleic Acids Res 35, 5262–5273 (2007).

29. Buck, J., Wacker, A., Warkentin, E., Wohnert, J., Wirmer-Bartoschek, J., Schwalbe, H. Influence of ground-state structure and Mg2+ binding on folding kinetics of the guanine-sensing riboswitch aptamer domain. Nucleic Acids Res 39, 9768–9778 (2011).

30. Hennelly, S. P., Novikova, I. V., Sanbonmatsu, K. Y. The expression platform and the aptamer: cooperativity between Mg2+ and ligand in the SAM-I riboswitch. Nucleic Acids Res 41, 1922–1935 (2013).

31. Suddala, K. C., Wang, J., Hou, Q., Walter, N. G. Mg(2+) shifts ligand-mediated folding of a riboswitch from induced-fit to conformational selection. J Am Chem Soc 137, 14075–14083 (2015).

32. Trausch, J. J., Marcano-Velazquez, J. G., Matyjasik, M. M., Batey, R. T. Metal Ion-Mediated Nucleobase Recognition by the ZTP Riboswitch. Chem Biol 22, 829–837 (2015).

33. Chen, B., LeBlanc, R., Dayie, T. K. SAM-II Riboswitch Samples at least Two Conformations in Solution in the Absence of Ligand: Implications for Recognition. Angew Chem Int Ed Engl 55, 2724–2727 (2016).

34. Xu, X., Egger, M., Chen, H., Bartosik, K., Micura, R., Ren, A. Insights into xanthine riboswitch structure and metal ion-mediated ligand recognition. Nucleic Acids Res 49, 7139–7153 (2021).

35. Ward, W. L., Plakos, K., DeRose, V. J. Nucleic acid catalysis: metals, nucleobases, and other cofactors. Chem Rev 114, 4318–4342 (2014).

36. Gonzalez, R. L., Jr., Tinoco, I., Jr. Identification and characterization of metal ion binding sites in RNA. Methods Enzymol 338, 421–443 (2001).

37. Zheng, H., Shabalin, I. G., Handing, K. B., Bujnicki, J. M., Minor, W. Magnesium-binding architectures in RNA crystal structures: validation, binding preferences, classification and motif detection. Nucleic Acids Res 43, 3789–3801 (2015).

38. Philips, A., Milanowska, K., Lach, G., Boniecki, M., Rother, K., Bujnicki, J. M. MetalionRNA: computational predictor of metal-binding sites in RNA structures. Bioinformatics 28, 198–205 (2012).

39. Sun, L. Z., Chen, S. J. Monte Carlo Tightly Bound Ion Model: Predicting Ion-Binding Properties of RNA with Ion Correlations and Fluctuations. J Chem Theory Comput 12, 3370–3381 (2016).

40. Sun, L. Z., Zhang, J. X., Chen, S. J. MCTBI: a web server for predicting metal ion effects in RNA structures. RNA 23, 1155–1165 (2017).

41. Zhou, Y., Chen, S.-J. Graph deep learning locates magnesium ions in RNA. QRB Discovery 3, e20 (2022).

42. Hayes, R. L., et al. Magnesium fluctuations modulate RNA dynamics in the SAM-I riboswitch. J Am Chem Soc 134, 12043–12053 (2012).

43. Cunha, R. A., Bussi, G. Unraveling Mg(2+)-RNA binding with atomistic molecular dynamics. RNA 23, 628–638 (2017).

44. Fischer, N. M., Poleto, M. D., Steuer, J., van der Spoel, D. Influence of Na+ and Mg2+ ions on RNA structures studied with molecular dynamics simulations. Nucleic Acids Res 46, 4872–4882 (2018).

45. Sponer, J., et al. RNA Structural Dynamics As Captured by Molecular Simulations: A Comprehensive Overview. Chem Rev 118, 4177–4338 (2018).

46. Hu, G., Zhou, H. X. Binding free energy decomposition and multiple unbinding paths of buried ligands in a PreQ1 riboswitch. PLoS Comput Biol 17, e1009603 (2021).

47. Sarkar, R., Jaiswar, A., Hennelly, S. P., Onuchic, J. N., Sanbonmatsu, K. Y., Roy, S. Chelated Magnesium Logic Gate Regulates Riboswitch Pseudoknot Formation. J Phys Chem B 125, 6479–6490 (2021).

48. Bao, L., Kang, W. B., Xiao, Y. Potential effects of metal ion induced two-state allostery on the regulatory mechanism of add adenine riboswitch. Commun Biol 5, 1120 (2022).

49. He, W., Henning-Knechtel, A., Kirmizialtin, S. Visualizing RNA Structures by SAXS-Driven MD Simulations. Front Bioinform 2, 781949 (2022).

50. Gohlke, H., Kiel, C., Case, D. A. Insights into protein-protein binding by binding free energy calculation and free energy decomposition for the Ras-Raf and Ras-RalGDS complexes. J Mol Biol 330, 891–913 (2003).

51. Case, D. A., et al. Amber 18. (ed^(eds). University of California: San Francisco (2018).

52. Jorgensen, W. L., Chandrasekhar, J., Madura, J. D., Impey, R. W., Klein, M. L. Comparison of Simple Potential Functions for Simulating Liquid Water. J Chem Phys 79, 926–935 (1983).

53. Cheatham, T. E., 3rd, Cieplak, P., Kollman, P. A. A modified version of the Cornell et al. force field with improved sugar pucker phases and helical repeat. J Biomol Struct Dyn 16, 845–862 (1999).

54. Perez, A., et al. Refinement of the AMBER force field for nucleic acids: improving the description of alpha/gamma conformers. Biophys J 92, 3817–3829 (2007).

55. Zgarbova, M., et al. Refinement of the Cornell et al. Nucleic Acids Force Field Based on Reference Quantum Chemical Calculations of Glycosidic Torsion Profiles. J Chem Theory Comput 7, 2886–2902 (2011).

56. Li, P., Roberts, B. P., Chakravorty, D. K., Merz, K. M., Jr. Rational Design of Particle Mesh Ewald Compatible Lennard-Jones Parameters for +2 Metal Cations in Explicit Solvent. J Chem Theory Comput 9, 2733–2748 (2013).

57. Joung, I. S., Cheatham, T. E., 3rd. Determination of alkali and halide monovalent ion parameters for use in explicitly solvated biomolecular simulations. J Phys Chem B 112, 9020–9041 (2008).

58. Frisch, M. J., et al. Gaussian 16 Rev. C.01. (ed^(eds) (2016).

59. Bayly, C. I., Cieplak, P., Cornell, W., Kollman, P. A. A well-behaved electrostatic potential based method using charge restraints for deriving atomic charges: the RESP model. J Phys Chem 97, 10269–10280 (2002).

60. Wang, J., Wolf, R. M., Caldwell, J. W., Kollman, P. A., Case, D. A. Development and testing of a general amber force field. J Comput Chem 25, 1157–1174 (2004).

61. Pastor, R. W., Brooks, B. R., Szabo, A. An analysis of the accuracy of Langevin and molecular dynamics algorithms. Molecular Physics 65, 1409–1419 (2006).

62. Berendsen, H. J. C., Postma, J. P. M., Vangunsteren, W. F., Dinola, A., Haak, J. R. Molecular-Dynamics with Coupling to an External Bath. J Chem Phys 81, 3684–3690 (1984).

63. Darden, T., York, D., Pedersen, L. Particle mesh ewald-and.Log(N) method for ewald sums in large systems. J Comput Phys 98, 10089–10092 (1993).

64. Ryckaert, J. P., Ciccotti, G., Berendsen, H. J. C. Numerical-Integration of Cartesian Equations of Motion of a System with Constraints - Molecular-Dynamics of N-Alkanes. J Comput Phys 23, 327–341 (1977).

65. Roe, D. R., Cheatham, T. E., 3rd. PTRAJ and CPPTRAJ: Software for Processing and Analysis of Molecular Dynamics Trajectory Data. J Chem Theory Comput 9, 3084–3095 (2013).

66. Humphrey, W., Dalke, A., Schulten, K. VMD: Visual molecular dynamics. J Mol Graph 14, 33–38 (1996).

67. Michaud-Agrawal, N., Denning, E. J., Woolf, T. B., Beckstein, O. MDAnalysis: a toolkit for the analysis of molecular dynamics simulations. J Comput Chem 32, 2319–2327 (2011).

