## Supplementary Table and Figures for "Magnesium ions mediate ligand binding and conformational transition of the SAM/SAH riboswitch"

Supplementary Materials

**Table S1.** Systems and molecular dynamics simulation lengths.

| Systems | # of $Mg^{2+}$ ions | Models | # of rep × MD length (μs) | Sum (μs) |
|---|---|---|---|---|
| SAH / SAM | 41 (Leap) | 10 | 4×1 = 4 | 80 |
| Apo | 41 (Leap) | 1 | 4×1 = 4 | 4 |
| SAH / SAM/ Apo | 25 (MCTBI) | 1 | 4×1 = 4 | 12 |
| SAH / SAM / Apo | 21 (Leap) | 1 | 4×1 = 4 | 12 |
| SAH / SAM/ Apo | 0 | 1 | 4×1 = 4 | 12 |

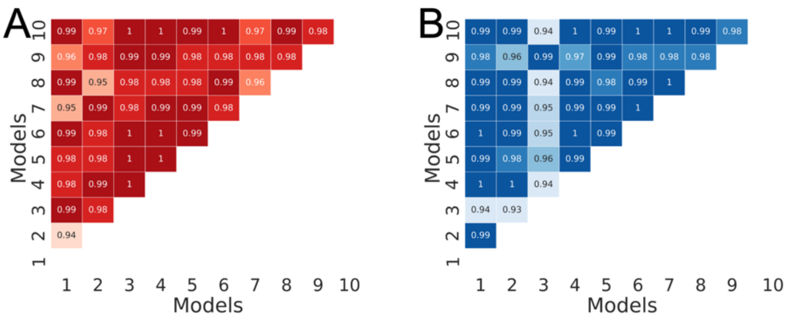


**Figure S1.** Correlations of nucleotide-ligand interaction energies between starting NMR models. (A) SAH. (B) SAM.

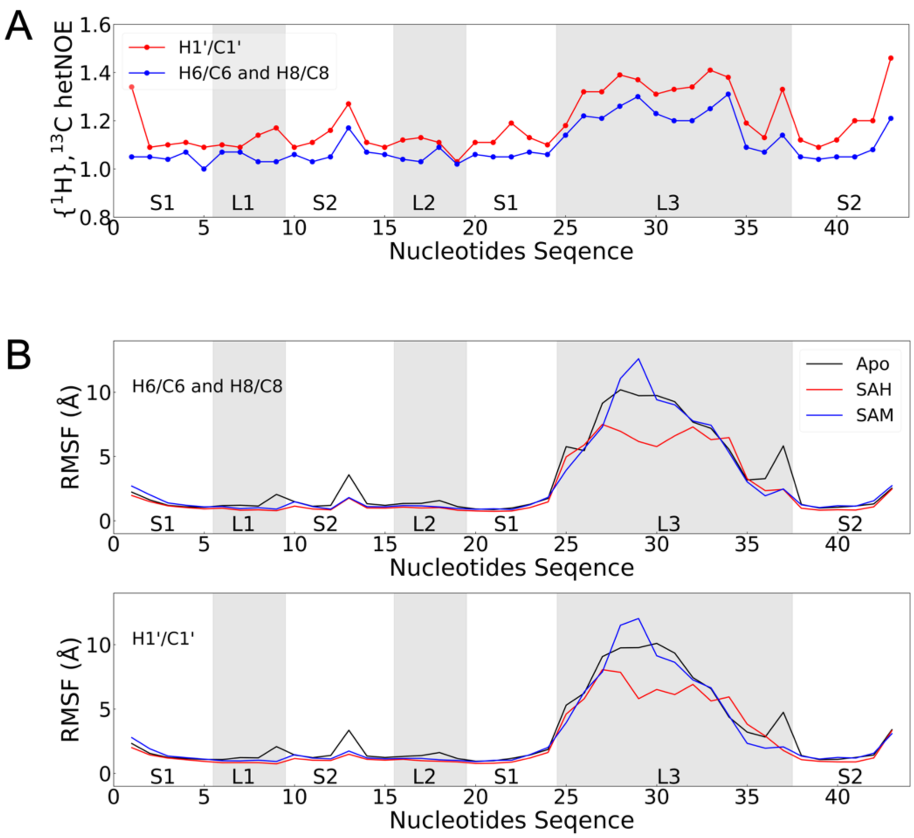


**Figure S2.** Flexibility profiles determined by NMR and MD simulations. (A) $^{1}$H-$^{13}$C heteronuclear Overhauser effects in the SAH-bound form, replotted using data reported by Weickhmann *et al.* [*Nucleic Acids Res* **47**, 2654-2665 (2019)]. (B) RMSFs from MD simulations.

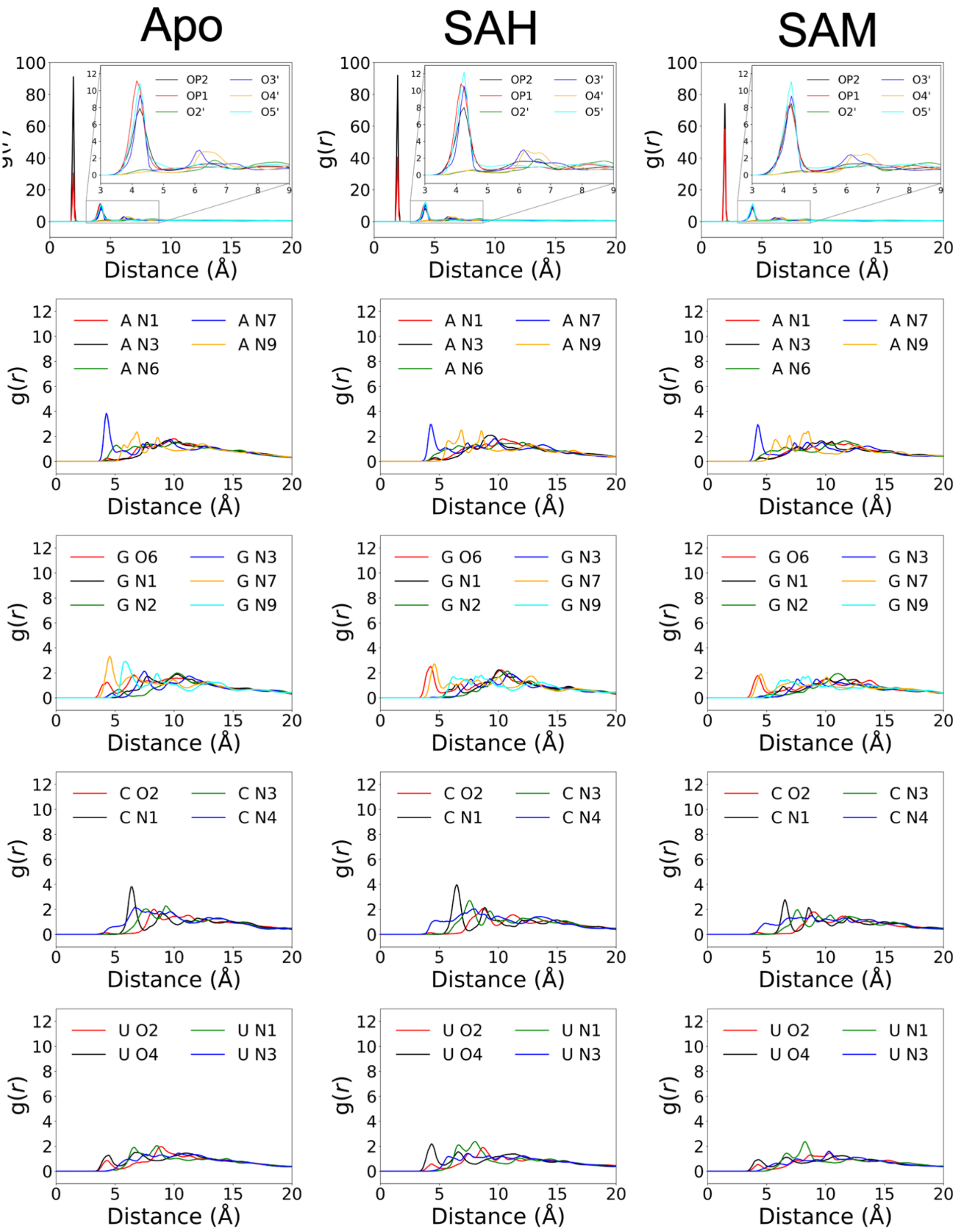


**Figure S3.** Radial distribution functions of $Mg^{2+}$ ions around backbone and base atoms in the apo and SAH and SAM-bound forms.

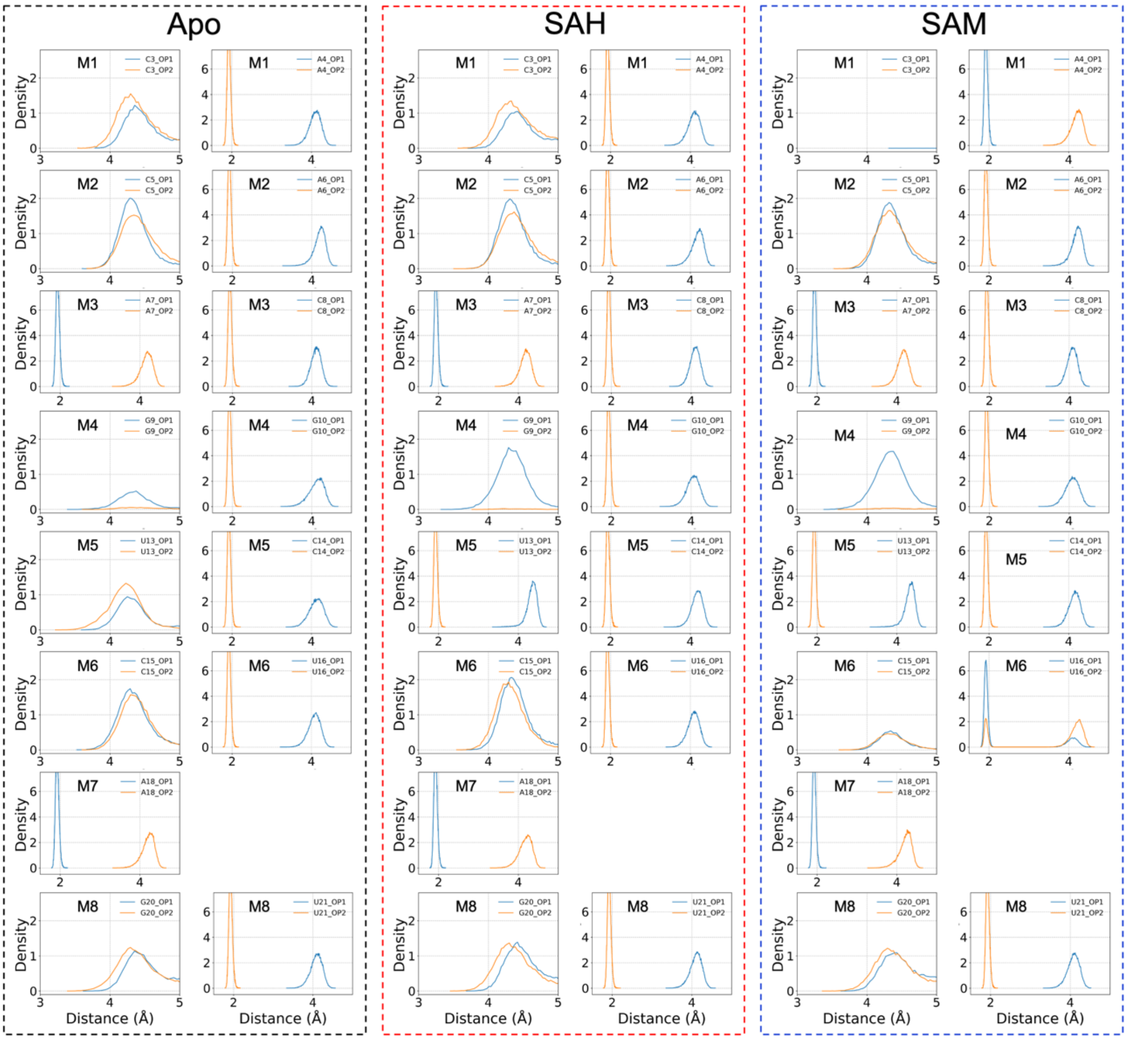


**Figure S4.** The distributions of distances between phosphate OP1 and OP2 atoms and conserved $Mg^{2+}$ ions.

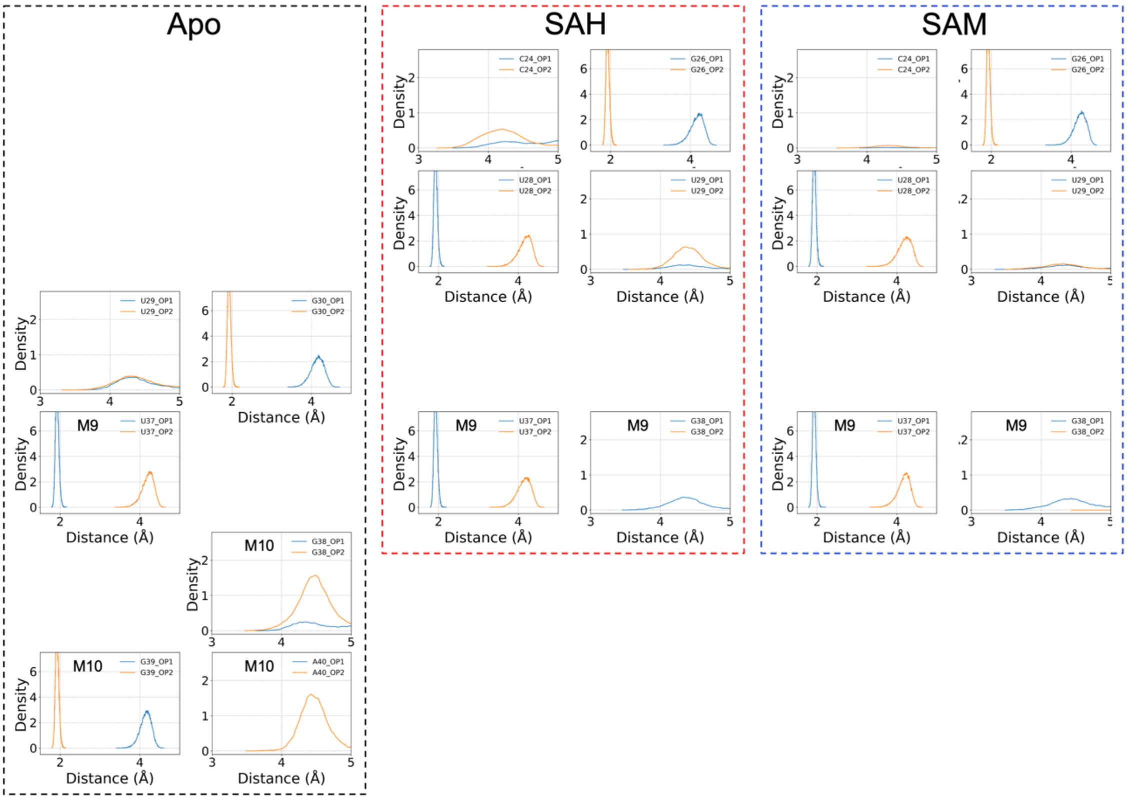


**Figure S5.** The distributions of distances between phosphate OP1 and OP2 atoms and $Mg^{2+}$ ions showing distinction between the apo and liganded forms.

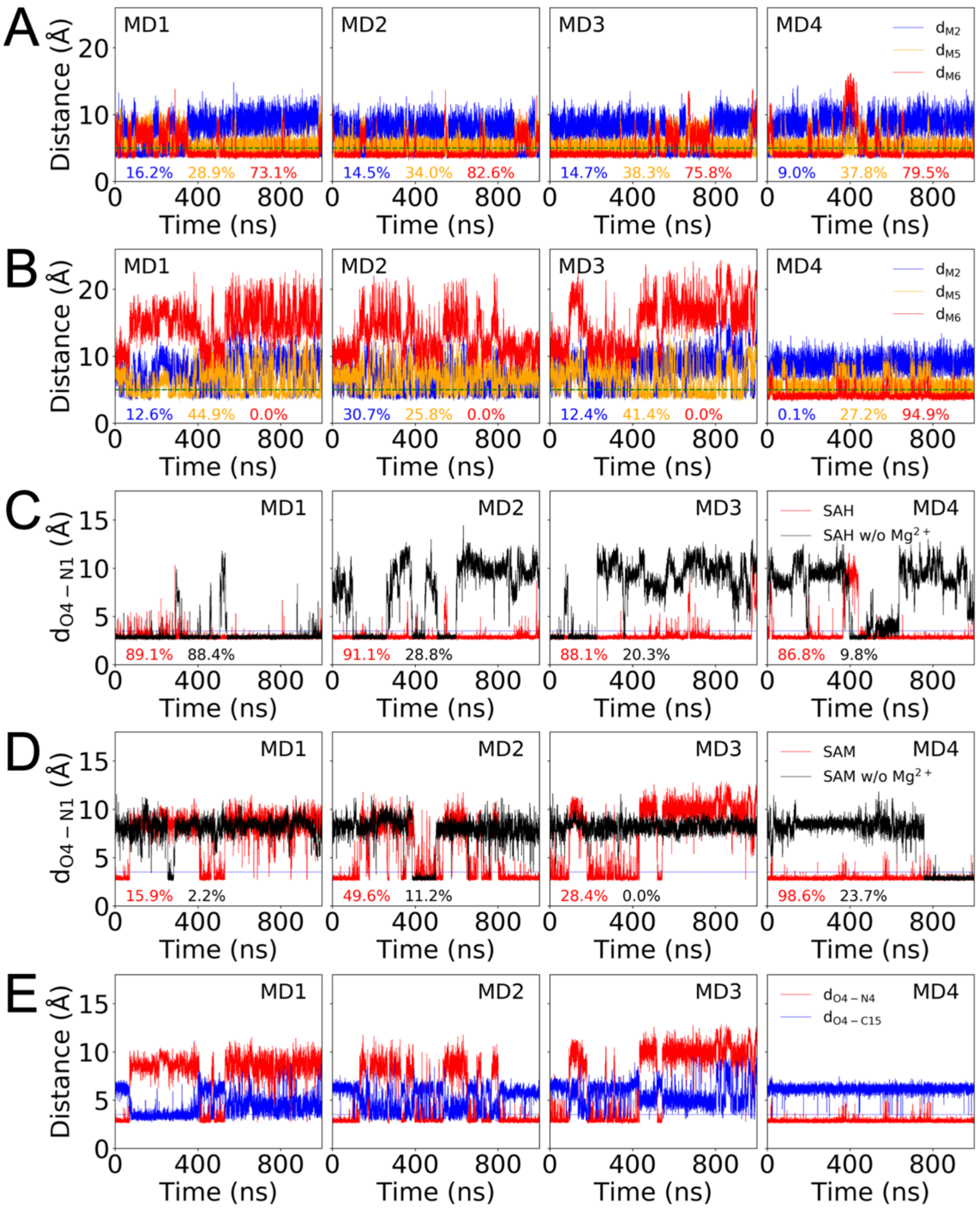


**Figure S6.** Traces of interatomic distances in four replicate simulations. (A) Distances of the SAH carboxy moiety to three inner-shell $Mg^{2+}$ ions. A horizontal line is drawn at 5 Å to indicate the cutoff for outer-shell coordination. The fractions of frames with outer-shell coordination are shown as percentages. (B) Corresponding results for the SAM-bound form. (C) The O4-N1 distances in the SAH-bound form without (labeled as "w/o") or with saturating $Mg^{2+}$. A horizontal line at 3.5 Å indicates the cutoff for hydrogen bond formation. The fractions of frames forming the hydrogen bond are shown as percentages. (D) Corresponding results for the SAM-bound form. (E) The O4-N1 and O4-C15 distances for the SAM-bound form with saturating $Mg^{2+}$.

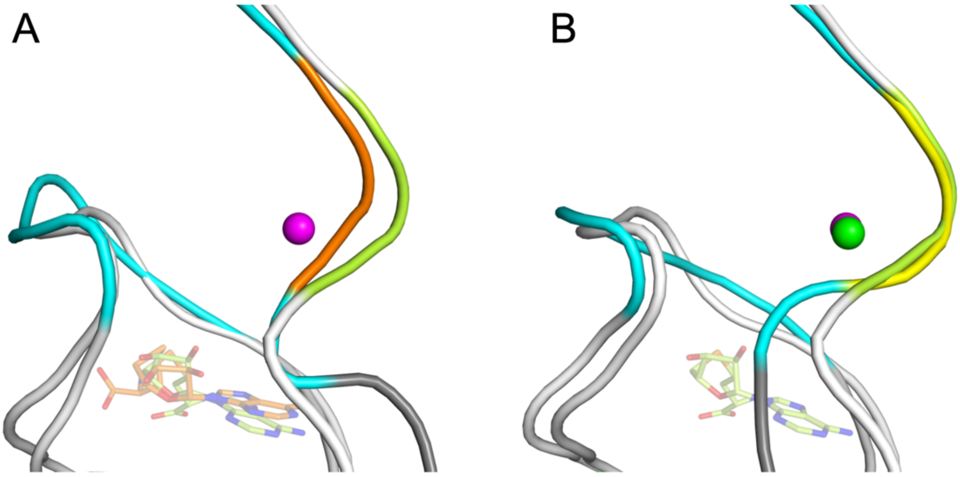


**Figure S7.** Comparison of the G39-A40-G41 backbone shapes between 6HAG, 6LY5, and a representative structure from the simulations of the apo form. (A) Superposition of 6HAG and 6LY5. G39-A40-G41 is shown in orange for 6HAG and limon for 6LY5; the ligands are shown with carbon atoms in the same two colors; note the different orientations of the carboxy moiety. A $Na^+$ ion from 6HAG is shown as a magenta sphere. (B) Similar comparison but 6HAG is replaced by the MD structure for the apo form and the color changed from orange to yellow. M10 is shown as a green sphere.
